# Current transcriptome database and biomarker discovery for immunotherapy by immune checkpoint blockade

**DOI:** 10.1101/2024.12.09.627506

**Authors:** Han Zhang, Binfeng Lu, Xinghua Lu, Anwaar Saeed, Lujia Chen

## Abstract

Immune checkpoint blockade (ICB) has revolutionized the current immuno-oncology and significantly improved clinical outcome for cancer treatment. Despite the advancement in clinics, only a small subset of patients derives immune response to the ICB therapy. Therefore, a robust predictive biomarker that identifies potential candidate becomes increasingly crucial in delivering this technology to the public. In this review, we first discuss the biomarkers that focus on tumor genome, tumor microenvironment and tumor-host interaction. Then, we compare existing databases for biomarker discovery for ICB response. We also present IOhub - an interactive web portal that incorporates 36 bulk and 10 single-cell transcriptome datasets for benchmark analysis of the current biomarkers. Finally, we highlight the trending interest in antibody drug conjugate and combination treatment and their use in precision immuno-oncology.

## Introduction

Immunotherapy by immune checkpoint blockades (ICBs) have significantly impacted clinical cancer treatment by targeting co-inhibitory ligand receptor pairs between immune cells and cancer cells, thereby reactivating the cancer-immunity cycle^1^. The humanized anti-cytotoxic T lymphocyte antigen 4 (CTLA4) antibody Ipilimumab, which is the first-approved ICB by the FDA in 2011, has remarkably doubled the 10-year survival rate for metastatic melanoma compared to historical data^2^. Following this, the blockade of other immune checkpoint molecules, programmed cell death 1 (PD1), or its ligand, PD1 ligand 1 (PDL1), has been demonstrated to offer a survival advantage across various malignancies, such as melanoma, non-small cell lung cancer (NSCLC), urothelial cancer and so on^3,4^; lymphocyte-activation gene 3 (LAG-3) has also been approved for use in advanced melanoma patients in combination with anti-PD1^5^. Compared with anti-CTLA4, anti-PD1/L1 or the combined treatment showed higher response rates and fewer side effects, resulting in the approval of ICBs as either second-line or first-line therapies for an expanding number of cancer types^6^.

Given the significant advancements in clinical care, one of the major challenges of current ICB therapy is that only a limited number of patients would respond to the treatment and derive clinical benefit^7^. Thus, there exists a strong interest in identifying and developing predictive biomarkers, both to enhance precision immuno-oncology and to understand the cancer resistance mechanisms^8^. Recent studies also highlight the necessity of predictive biomarker and use of single cell transcriptome in helping patient stratification^9,10^. For example, colorectal cancer patient with high microsatellite instability showed significant overall response rate whereas patients with microsatellite stability are mostly non-responsive to the ICB, partially due to the profile of neutrophil^11^. Besides, some biomarkers fail to exhibit consistent efficacy across several studies in the same cancer type^8,12^.

To investigate the characteristics of the biomarkers for ICB, several databases and web portals have been released^13-15^. These databases have collected large amount of bulk transcriptome data of the tumor samples in ICB studies and carried out benchmark analysis of the transcriptome-based biomarkers across different cancer types. However, single-cell transcriptomic data of the responsive samples to the ICB are not considered into the analysis; moreover, the number of collected datasets needs to update frequently as immuno-oncology is becoming one of the hottest area in cancer therapy. In this review, we first focus on the current predictive biomarkers for ICB response and discuss the mechanisms underlying the interaction within the tumor microenvironment (TME); then, we present IOhub, an interactive web portal that tracks the latest transcriptome data of ICB-treated tumor samples. IOhub is consist of 36 bulk and 10 single-cell transcriptome datasets, making it the biggest database and web portal for ICB research. Finally, we highlight the cutting-edge advances in immuno-oncology, focusing on antibody-drug conjugate (ADC) and combination therapies.

### Current biomarkers for immune checkpoint blockade

Generally, there are three main strategies of biomarker discovery for ICB response prediction: 1) to estimate neo-antigen production ability from tumor genome; 2) to explore whether there exist enough dysfunctional T cells or tumor cells that are expressing PD1/L1 in the TME; 3) to look into the host profile and tumor-host interaction (**Table 1**).

### Biomarkers targeting tumor genome

The cancer cell is a dynamic entity that is characterized by millions of mutations. However, not all kinds of mutations have the capability to generate novel epitopes - specific part of the antigens that bind to the major histocompatibility complex (MHC). Consequently, it is logical that the total number of non-synonymous single nucleotide variables (nsSNV) in the tumor genome influences the likelihood of successful immune response by generating more immunogenic peptides so that it becomes easier for T cells to recognize the tumor cells^16^. One of the popular method of estimating tumor mutation burden (TMB) is to take the total number of nsSNV or missense variants in the tumor genome. The first significant positive correlation between TMB and immune response was found in patients treated with anti-CTLA-4 in melanoma study^17^. The finding was later validated in several other cancer types with anti-PD1 blockades^18,19^. Notably, some of the best ICI response rates were observed in carcinogen-driven cancers (i.e. melanoma and NSCLC) which typically have high mutation burdens because of tobacco use and ultraviolet light^20^. To date, researchers and clinics employ whole genome sequencing, whole exome sequencing or targeted sequencing to assess the TMB value via a couple of different pipelines, such as MSK-IMPACT panel^21^ and FoundationOne CDx panel^22^.

Despite its promise, TMB is not universally accepted as a predictive biomarker across all cancer types. The association between TMB level and clinical response is still unclear. For example, patients with clear-cell renal cell carcinoma showed no significant association between TMB level and response from inhibition of the PD1 pathway^23^. Additionally, in cancers such as metastatic urothelial cancer and melanoma, where a high TMB correlates with immune response^24,25^, TMB fails to reliably discriminate all responders from non-responders. Some patients with relatively low level TMB are still responding to the ICB treatment. Currently, there is no optimal cutoff for TMB level binary classification, indicating more effort and data are needed in identifying cancer type specific standard for this biomarker^8^.

Besides TMB, somatic copy number alteration (SCNA) could be another source of neo-antigen production^26^. In a review analysis of TCGA data, focal level CNV (covering < 50% chromosome arms) was found to sustain proliferation, implying an abnormal function of specific genes targeted by the focal CNVs; on the other hand, arm level CNV (covering >= 50% chromosome arms) was negatively associated with immune infiltration, suggesting mechanism related to general gene dosage imbalance rather than specific genes. The author also proposed an integrated biomarker that combined arm level CNV with mutation burden. In two different validation datasets of previous melanoma clinical trials, the integrated biomarker showed a better performance in predicting survival rate than using either mutation burden or CNV level alone, indicating arm level CNV would provide additional information independent of mutations. It might be because CNV contributes to the loss of genes that are needed for neoantigen presenting so that the T cells fail to identify the tumor cells. However, an evaluation in terms of SCNA performance in pan-cancer study is still needed to be validated in clinical trials.

Intragenic rearrangement (IGR) burden is type of mutation that emphasize the structural variation level within the coding genes^27^. In the analysis of pan-cancer, IGR was demonstrated significantly associated with immune infiltration in breast, ovarian, esophageal, and cervical cancers; it was also shown to be the most significant predictor against tumor infiltering lymphocyte on top of other genomic biomarkers in above cancer types. Further analysis in the IMVigor210 trial evaluating atezolizumab in advanced urothelial carcinoma shows that IGR burden increased in patients who had been treated with platinum-based chemotherapy and predicted clinical benefit among TMB-low, platinum-exposed patients. Overall, IGR suggests a new source of neoantigen and thus could serve as an important biomarker when the mutation burden is low. Similar to IGR, the frameshift indel burden also correlates with inflammatory signature and has the potential to predict ICB response by producing neoantigens in a subset of cancer types, such as head and neck squamous cell carcinoma (HNSCC)^28^ and renal cell carcinoma (RCC)^29^.

### Tumor microenvironment

While the biomarkers based on tumor genome focus on the ability of neoepitopes production, the biomarkers for tumor microenvironment will investigate two other attributes: 1) up regulation of either PD-1 expression on immune cells or PD-L1 expression on tumor cells; 2) immune phenotype - an inflamed tumor is more favorable of recruiting T cells into the microenvironment.

The primary mechanism of immune checkpoint blockade is to reinvigorate the dysfunctional T cells that are already present in the TME^30^. Therefore, the PD1/PDL1 expression is widely considered to be an essential factor in predicting immune response. Accordingly, the PD-L1 protein expression by immunohistochemistry (IHC) assay has been approved by FDA for several cancer types (e.g. NSCLC and triple negative breast cancer)^31^. The underlying rationale is that tumors with up-regulated PD-L1 expression is more likely evade immune detection by exploiting the PD-1/PD-L1 axis; thus, by blocking the interaction between PD-1 and PD-L1, it is assumed that the dysfunctional T cells inside the TME will be reactivated. Interestingly, the association between high PD-L1 expression and better immune response is inconsistent. In a Phase I trial nivolumab, the overall response rates of patients with advanced melanoma were found significantly higher in PDL1-positive group (Combined Positive Score >= 5%, suggesting over than 5% of PD-L1 staining cells divided by the total number of viable tumor cells)^32^. However, researchers failed to observe the same conclusion in subsequent trails^33^. Plausible reasons for these discrepancies include variability in IHC assays and inconsistent cutoff for PD-L1 positiveness (i.e., percentage of stained cells or staining intensity). In summary, the utility and criteria of using PD-L1 expression by IHC assay as a universal predictive biomarker for all cancers remain undefined^8^.

Similar to the PD-L1 expression, immune phenotype also reflects the presense of dysfunctional T cells within the TME. Based on the spatial distribution of CD8^+^ T cells in the microenvironment, three major immune phenotypes have been identified: inflamed, where immune cells infiltrate deeply into the TME; excluded, where immune cells accumulate while not penetrate into the tumor site efficiently due to transformation of the tumor associated fibroblast and extracellular matrix; desert, where immune cells are almost absent from the TME^34^. Inflamed tumors are thought to harbor pro-inflammatory cytokines, such as IL-2^35^ (interleukin-2, a key factor for T cell activation) and IFN*γ* (interferon gamma, which enhances the anti-tumor effect of CD8^+^ T cells). These cytokines not only attract the cytotoxic T lymphocytes coming out of blood vessel but also create an optimal place for T cell activation and expansion. Thus, more T cells would be reinvigorated after the blockade of PD1/PDL1 ligation in the inflamed tumors. However, the casual relationship between presence of pro-inflammatory cytokines in the TME and infiltrated CD8^+^ T lymphocytes^1^ is still unclear. Currently, histological analysis is widely regarded as the gold standard for phenotyping in clinics^36^. Some research teams tried to use binomial logistic LASSO (least absolute shrinkage and selection operator) based on gene expression data to predict the spatial distribution of CD8^+^ T cell; nevertheless, the model has shown limited accuracy in distinguishing different immune phenotypes^37^.

Due to the high cost of whole genome sequencing and whole exome sequencing, transcriptomic data becomes increasingly accessible in companion with patient’s clinical information. Gene expression biomarkers such like Oncotype DX^38^ (a profiling test that analyzes 21 gene expressions so as to predict how likely the patients will benefit from getting chemotherapy or hormone therapy after surgery) has demonstrated its efficacy in breast cancer. It is also hypothesized that gene expression signatures could provide valuable insight in predicting ICB response. A common approach for using gene expression as a biomarker is to assess the average expression level across a certain gene list. For example, the average gene expression of pro-inflammatory genes can indicate the degree of inflammation within the tumor^23,26,39^. Since the tumor microenvironment is a mixture of tumor cells, immune cells and normal cells, researchers often utilize the reference-based algorithms to deconvolute the relative proportions of different immune cell types in the tumor mRNA^40,41^. Recent deconvolution studies have also incorporated single-cell transcriptome data as the reference to infer the percentage of immune cell types more accurately; this has been demonstrated to generate more reliable estimation^42,43^.

### Gut microbiota

The intestinal microbiome has currently emerged as another promising biomarker for ICB response prediction since its extensive interactions with the host immune system^44^. Specific microbial species, for example *Bacteroides, Bifidobacterium* and *Akkermansia muciniphila*, have been reported in association with clinical benefit of ICB therapies^45^. Such microbes play a critical role in tumor-host immunity by promoting T cell activation and regulating cytokine production; thereby, facilitating immune cell infiltration into the TME. In fact, the abundance of these microbiome species has been found significantly higher in melanoma and RCC patients who experience clinical benefits compared to those who do not^45,46^. Previous study also implies that it is the composition of microbiome rather than any single species of bacteria that determines ICB response^47^. However, the diversity of microbiome largely depends on the tissue, environmental condition and prior antibiotic use, resulting in inconsistent result in large-scale analysis^44^. Despite these challenges, the findings highlight that the gut microbiota is becoming a significant factor in antitumor immunity and ICB response. In addition, the microbiota’s modifiable nature offers opportunities for intervention through dietary changes to improve patient clinical outcomes. Overall, further research is needed to standardize and integrate gut microbiome data with other biomarkers for more precise ICB response predictions.

### Current public databases for immune checkpoint blockade

To explore the spectrum of biomarkers for ICB response prediction, various databases and web portals have been developed^13-15,48^. These resources have collected bulk transcriptome data, overall survival, biopsy timepoint, immune response as well as other clinical information for survival analysis, deconvolution in the tumor microenvironment, differentially expressed gene analysis and so on. However, none of the current database has used the information from single-cell transcriptome of the responders to guide the analysis in bulk transcriptome data and has not provided the consistency test across studies in the same cancer type (**Table 2**).

### ICBAtlas

ICBAtlas is a well-organized database that is designed to analyze the transcriptomic biomarkers of ICB therapies across multiple cancer types^13^. Transcriptomic data and clinical information of 1,515 ICI-treated patient samples across 9 cancer types from 25 public databases were collected in this resource. The author further conducted analyses to compare gene expression signatures, differential expressed genes and GO terms between responders and non-responders, as well as between pre-treatment and on-treatment groups. The authors also introduced a Response Score, which measures the impact of indicative genes on ICB response, allowing researchers to identify potential gene-based biomarkers. Additionally, they examined immune cell infiltration through deconvolution in the tumor microenvironment and identified immune cell subpopulations that are associated with better clinical outcomes in different cancer types. The ICBAtlas provides users with the ability to explore these features at the gene, cancer type, or immune checkpoint level, offering a robust tool for understanding the molecular mechanisms behind ICB treatment response. The platform is freely accessible and aims to support researchers in identifying biomarkers and improving the efficacy of personalized cancer immunotherapy.

### TIDE

The TIDE score is a computation method that integrates signatures of both tumor immune dysfunction and exclusion to predict ICB response^48^. This method overcomes the limited sample size of immunotherapy by employing 33,197 non-ICB treated bulk samples from 189 previous human cancer studies. Instead of estimating the association between certain gene sets and immune response, the author calculated the interactions between each gene and cytotoxic T cell level in the single variate Cox-PH survival model. The normalized score from Wald test was then used to predict if the patient would respond to the anti-PDL1 treatment. To test the model, the author used publicly available data of melanoma patients after checkpoint therapy and reported the best AUROC (area under receiver operating characteristic curve) over other published biomarkers. Overall, this analysis provides insights into using the public databases to estimate the immune response; one limitation is that this method only works for melanoma and non-small cell lung cancer. Their website also supports the analysis of TIDE score and provides access to public transcriptome datasets.

### ICB-Portal

ICB-Portal is a comprehensive resource that provides benchmark results and detailed analysis of the existing biomarkers for ICB response^14^. It also offers an online platform on which researchers are able to test their custom biomarkers for predicting ICB response and clinical outcomes. This study provides valuable insights into the variability and performance of biomarkers. To identify robust biomarkers, the authors curated 29 transcriptomic datasets, including over 1,400 patients treated with ICB therapies in 5 cancer types. The researchers estimated 39 transcriptomic biomarkers and 48 scoring systems from canonical gene sets and deconvolution outcomes. These biomarkers were systematically evaluated to determine their predictive efficacy for ICB response, overall survival as well as progression-free survival across different datasets, cancer types, biopsy times and treatment arms. Among all available biomarkers, TIDE and cytotoxicity score, showed the best predictive performance for ICB response. However, one of the major limitation is the small number of cancer types that are considered. The majority of the datasets collected in this resource are from melanoma, NSCLC and metastatic urothelial cancer, which has been proved highly correlated with cytotoxic and inflammatory signatures as a whole. Therefore, more efforts are needed to extend the biomarker evaluation to other cancer types.

### TIGER

Compared with other existing databases and web portals, TIGER (Tumor Immunotherapy Gene Expression Resource) has advantages in the collection of single-cell transcriptome data and providing extensive analytical modules for ICB response prediction^15^. Collectively, the author curated over than 1,500 tumor samples with clinical outcomes from 18 bulk transcriptome datasets. To show the full landscape of immune cell subpopulation, this resource also encompasses single-cell sequencing data for over 2.1 million immune cells from 655 samples. TIGER supports a variety of features for users to explore the associated pathways, trajectory mapping and response prediction for the inquired data. Despite all these features, one of the concerns is that the AUROC analysis for each biomarker is assessed on both pre-ICB treated and post-ICB treated tumor samples as the whole population; this could introduce bias and overfitting of the metric evaluation as the batch effect of the biopsy time and variance within the same patients are taken into the calculation.

### IOhub

As shown, the current databases have not taken advantages of the single-cell transcriptome which offers more insights at individual cell and subtype; besides, they rarely considered the variability of the biomarker across different studies in the same cancer type. To solve this limitation, we present IOhub, an interactive data repository that tracks the publicly available IO treatment transcriptomic data (**Figure 1A**) that integrates 36 bulk and 10 single-cell datasets to indicate the immune response to ICB (**Table 3**).

**Figure 1.**
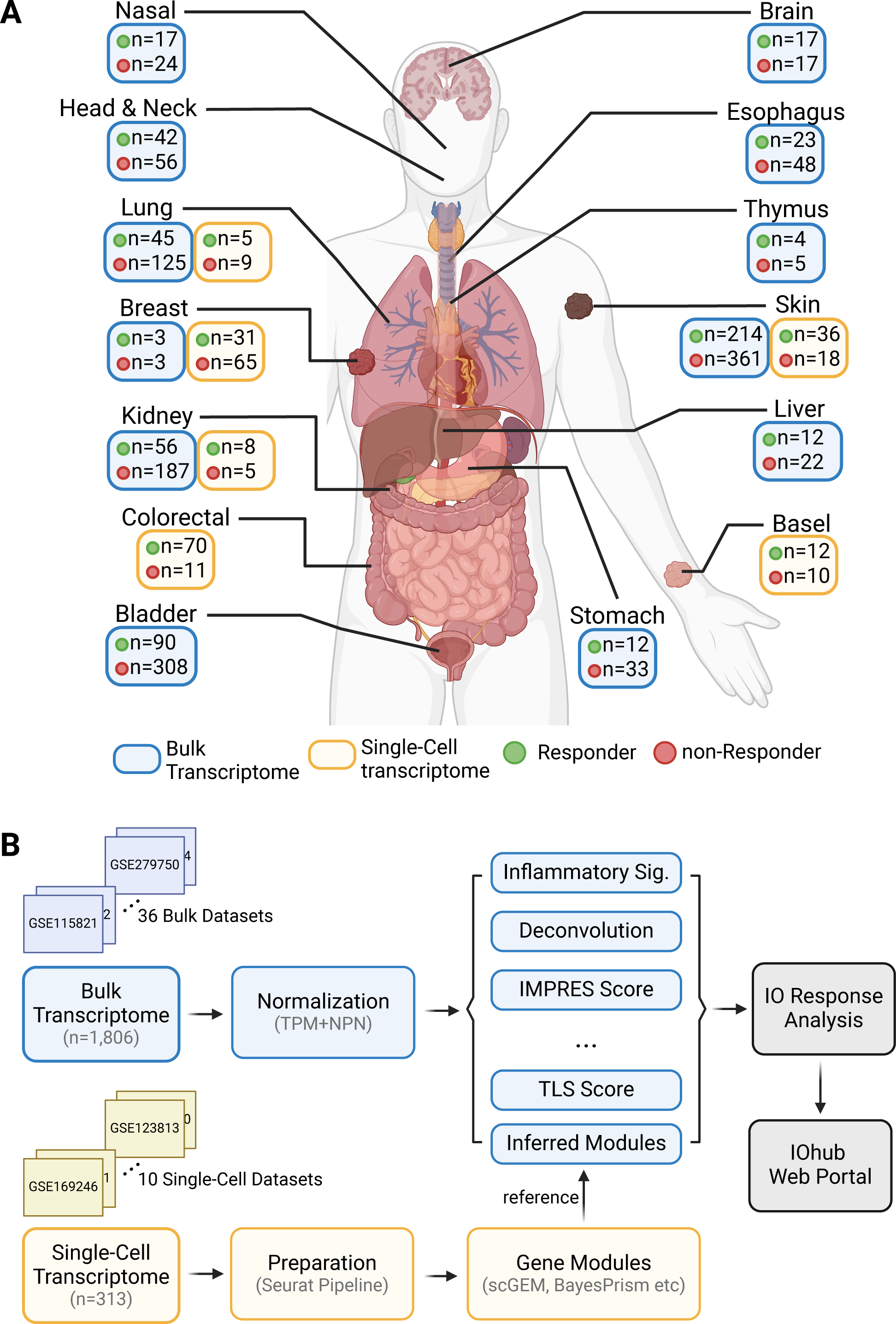
Overview of IOhub. (**A**) An illustration of IOhub database. The bulk transcriptome data is labeled in blue box and single-cell transcriptome data is labeled in yellow box. Responsive tumor samples and non-responsive tumor samples are colored in green and red respectively. All cancer types are directed to corresponding organs. (**B**) Workflow of analysis in IOhub. Processes in blue boxes and yellow are affiliated with bulk and single-cell transcriptome analysis respectively. The normalization of bulk data is done separately in each dataset.

Transcriptomic data as well as clinical information were obtained using the public dataset IDs. Patients are labeled as “Response/R” if they showed clinical benefit, complete response (CR), partial response (PR), mixed response (MR) or clonal expansion after ICB treatment; patients are labeled as “non-Response/NR” if they showed no clinical benefit, progressive disease (PD), stable disease (SD) or no clonal expansion after ICB treatment. CR/PD/SD/PD is determined based on RECIST v1.1. For each bulk-RNAseq dataset, gene expressions were normalized in transcript per million (TPM) formats. We then used non-paranormal normalization to remove batch effects and covariates of cancer type (**Figure 1B**). Samples from CM-009 cohort in Braun’s study were removed as they are duplicated in Miao’s study. In total, 1806 ICB treated tumor samples are included from 36 datasets, consisting of 1496 pre-treatment samples, 310 post-treatment samples, 535 responsive samples, 1189 non-responsive samples and 82 non-evaluated samples in 14 cancer types (**Figure 2A**). As for scRNAseq datasets, we extracted the raw single cells and further filtered if 1) the number of genes is less than 250 or higher than 8000; 2) the mitochondrial ratio is greater than 20%; 3) the number of unique molecular identifier (UMI) counts is less than 500. To rule out the bias from manual annotation, we used SingleR to map the cell major type and subtype to Monaco immune database. In total, we collected 593888 responsive and 280564 non-responsive single cells from 162 and 118 samples respectively in 7 cancer types (**Figure 2B**). A two-round clustering using the top 10 to 20 principal components and resolution of Louvain community detection ranging from 0.4 to 1.0, depending on the visual clustering performance in UMAP plot, was applied for each single cell cohort. The processed single cell expressions were feed into the Bayesian models scGEM^43^ and BayesPrism^42^ to learn gene co-expressing modules for bulk deconvolution.

**Figure 2.**
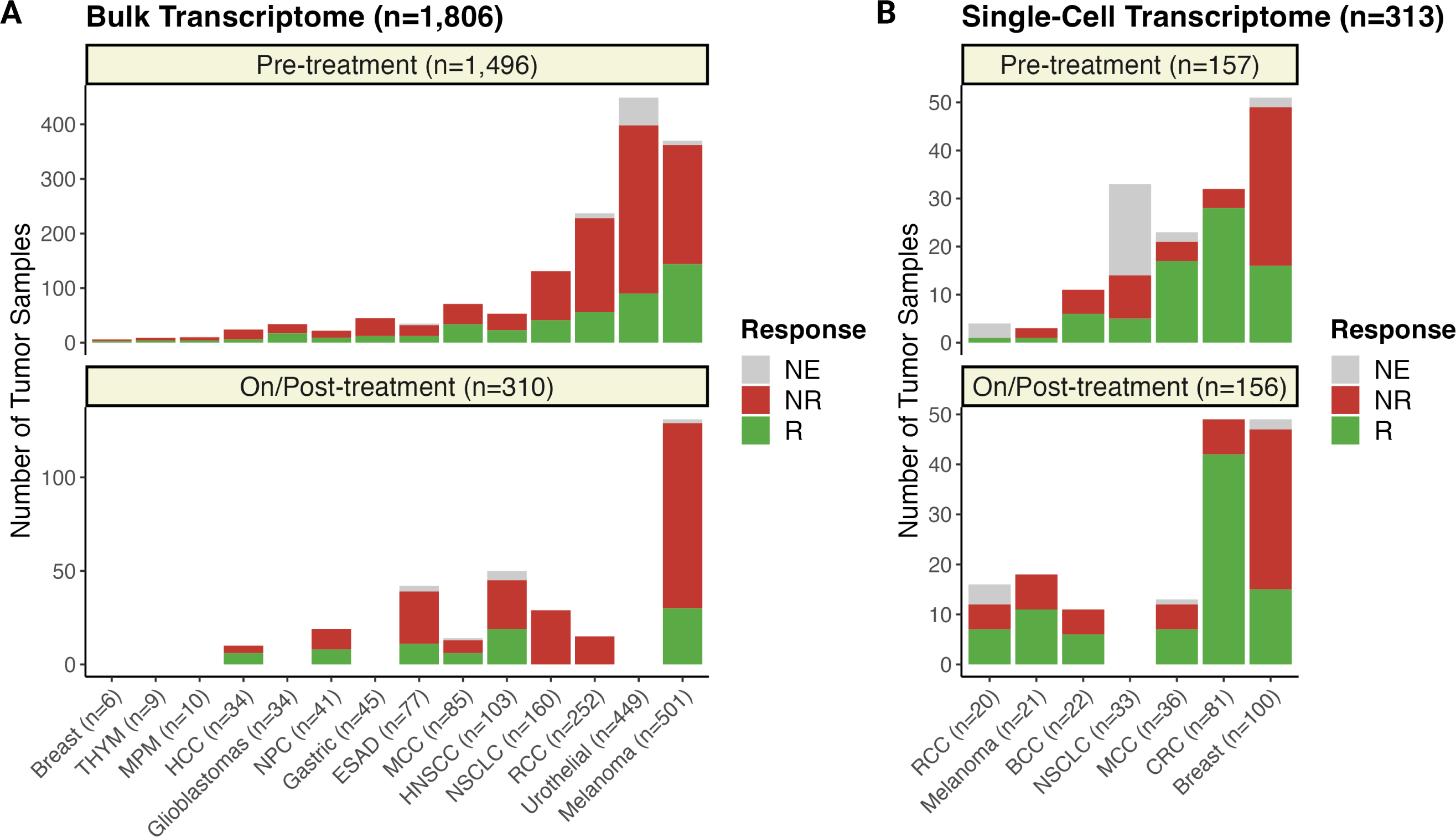
Sample distribution in IOhub. (**A**) The distribution of tumor samples in bulk transcriptome by cancer type. The barplot is ranked by the ascending order of the total sample size within each cancer type. Responsive, non-responsive and non-evaluated tumor samples are colored in green, red and grey respectively. The barplot is split by the biopsy timepoint (Pre-ICB and On/Post-ICB). (**B**) The distribution of tumor samples in single-cell transcriptome by cancer type. The total sample sizes are provided.

Interestingly, we found that a number of the transcriptomic biomarkers are predictive of immune response to ICB in melanoma (**Figure 3A**). Microsatellite instability score is significantly higher in responder group in gastric and esophageal cancer, which is consistent with previous studies^31,49^. Of note, no universal transcriptomic biomarker for pan-cancer was found. Even for informative biomarkers in the melanoma, they fail to be applicable to all study cohorts (**Figure 3B**). As is shown, the IFN*γ* score is significantly higher in study cohorts of Augustin, Gide and Prat with Wilcox p-values of 0.03, 3e-07 and 0.042 respectively; while T cell gamma delta, calculated from Cibersort output, correlates with ICB response in Gide, Jaiswal and Riaz with Wilcox p-values of 0.007, 0.008 and 0.003 respectively. Overall, it reveals the considerable variance of biomarker efficacy within the same cancer type, implying the benchmark analysis of an effective biomarker should be evaluated across all cohorts rather than look into the total population of melanoma tumor samples. Finally, we employed the single cell transcriptomic data of ICB-treated samples to discover new biomarkers for RCC tumor samples as almost all current transcriptomic biomarkers are not associated with ICB response in kidney cancer (**Figure 3A**). Two scGEM model were trained using Bi (around 8,000 cells) and Krishna (around 50,000 cells) single cell datasets. We observed that as the number of single cells increases in the model, the Wilcox p-value between responder and non-responder is getting more significant for the same deconvoluted gene co-expressing module in two models. Compared with other existing biomarkers that relate to the extracellular matrix and stromal program and IFN*γ* score, the single-cell based biomarker from scGEM shows clearer trend and significant difference between responsive tumors and non-responsive tumors in RCC (**Figure 3C**). In summary, the single-cell referenced based deconvolution method will offer more opportunities in biomarker discovery as the model sees more cells in the future.

**Figure 3.**
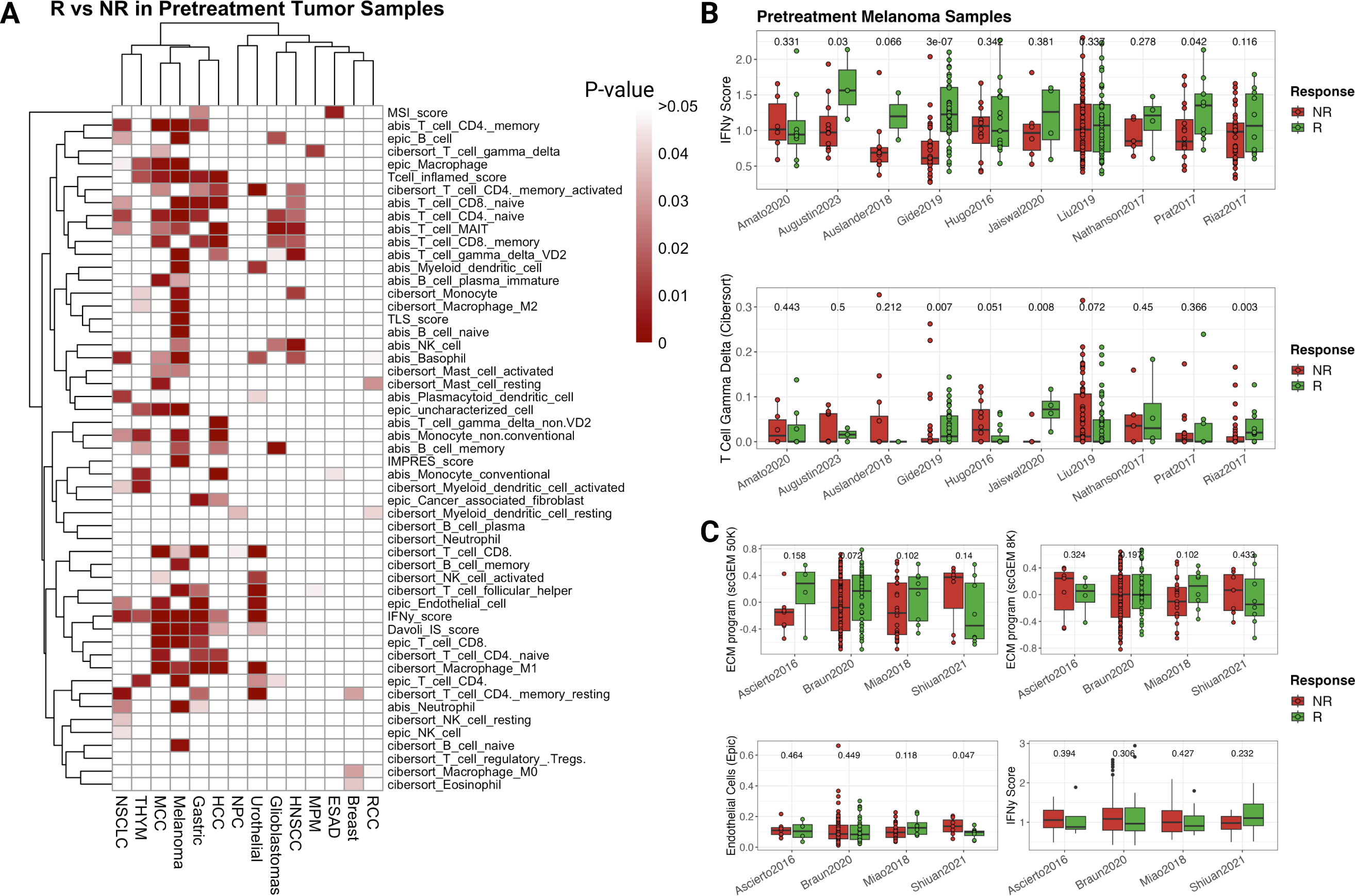
Transcriptome biomarkers show distinct correlations with ICB response across cancer types and studies. (**A**) Heatmap of the p-values of one-sided Wilcox test for responsive tumor samples against non-responsive tumor samples in corresponding biomarkers. P-values over than 0.05 are colored in white; p-values equal to 0 are colored in dark red. (**B**) Boxplots with jitter points of the IFN*γ* score and T cell gamma delta score that is estimated by Cibersort in 10 melanoma studies. (**C**) Boxplots with jitter points of ECM program scores (estimated by scGEM with 50,000 and 8,000 RCC ICB-treated single cells), the IFN*γ* score and endothelial score that is estimated by Epic in 4 RCC studies. The p-values shown on top of each boxplot represent the exact p-value of two-sided Wilcox test for responder against non-responder samples. Only pre-treatment tumor samples are used.

To assist the exploration of biomarker discovery in ICB response prediction, we also provide a R Shiny App (https://shiny.crc.pitt.edu/iohub/) that allows users to compare their interested biomarker and gene distribution across all datasets (**Figure 4A, B**). More features such as overall survival analysis and differentially expressed genes as well as the detailed benchmark evaluation across cohorts will be released in the next version soon.

**Figure 4.**
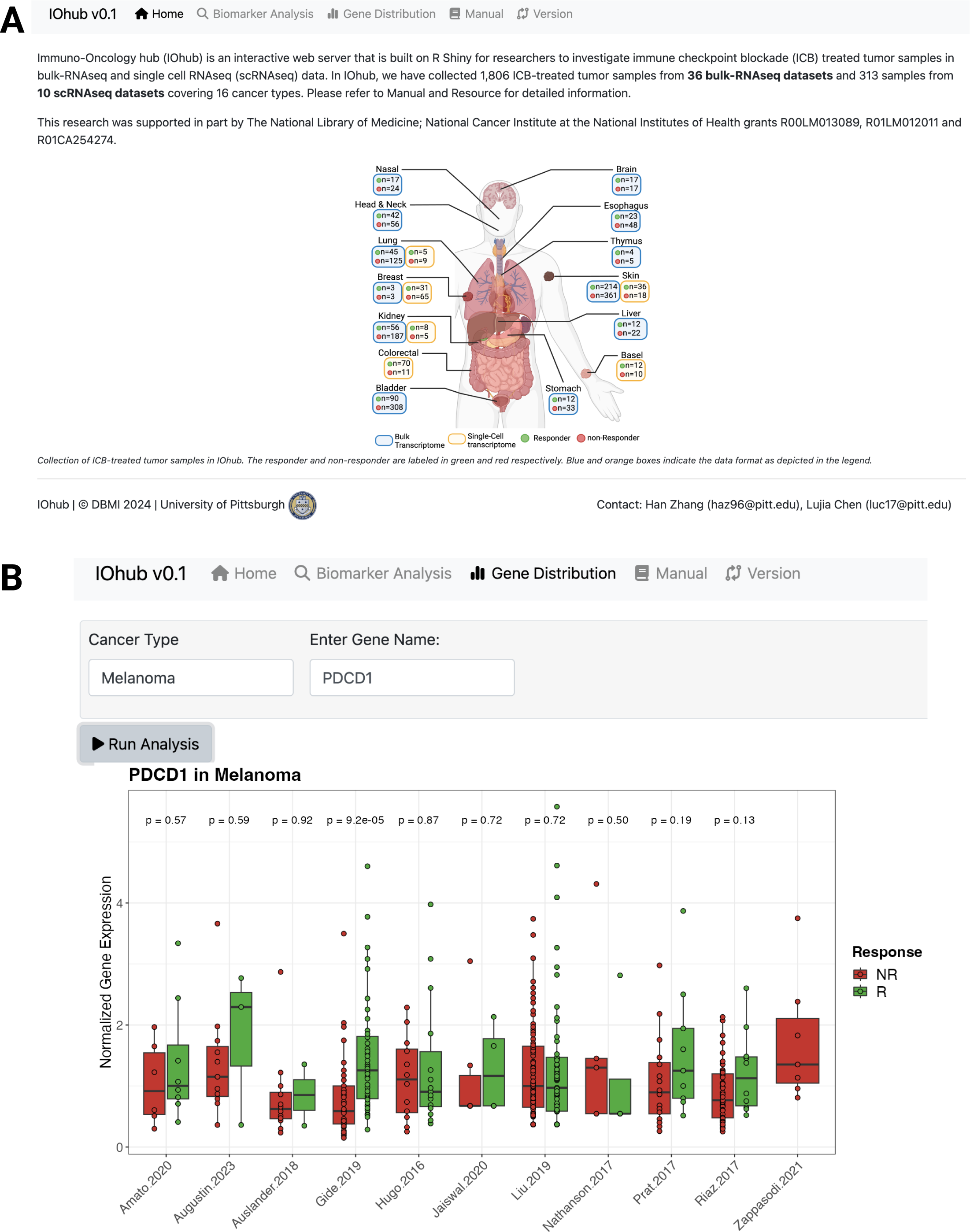
Illustration of IOhub web portal. (**A**) The home page of IOhub describing the purpose and funding of this study. (**B**) View of the feature of exploring gene expression distributions across different curated studies of the selected cancer type. Boxplots with jitter points of the responsive and non-responsive tumor samples are colored in green and red respectively. The p-values shown on top of each boxplot represent the exact p-value of two-sided Wilcox test for responder against non-responder samples. Only pre-treatment tumor samples are used.

### Future direction

ICB has revolutionized the treatment of cancers by harnessing the host immune system to recognize and eradicate tumor cells. However, a significant challenge remains – the low response rate and potential relapse to the ICB treatment^31^. To address these limitations, future directions in immunotherapy are focused on improving the overall response rate and survival outcomes through innovative approaches such as the antibody drug conjugate (ADC) and combination treatment to enable precision immuno-oncology.

ADCs have attracted much attention in the past five year as a transformative modality in immuno-oncology. These molecules comprise three key components: a monoclonal antibody that selectively binds to antigens expressed on tumor cells, a linker that connects the antibody to the cytotoxic payload and the payload, which is typically a highly potent chemotherapy agent^50^. Therefore, it offers targeted delivery of potent cytotoxic agents directly to tumor cells. The effector mechanism begins with the ADC binding to its target antigen on the tumor cell surface, leading to internalization of the ADC-antigen complex. Once internalized, the linker releases the cytotoxic payload within the tumor cell in lysosomes, where it induces cell death by disrupting critical cellular processes, such as microtubule formation or DNA replication. After the death of tumor cell, the payload will also penetrate into adjacent tumor cells to trigger cell death using the same mechanism (bystander effect)^51^. In recent days, the industry has shown growing interest in bispecific ADC (bsADC)^52^. Compared with ADC, bsADC targets two tumor associated antigens simultaneously; thereby exhibits improved efficacy due to synergistic effects, increased internalization rate and overcomes tumor heterogeneity, reducing resistance and off-target side effects. With the improvements in payload potency, drug-to-antibody ratio, linker stability, and target specificity, ADCs are addressing the unmet clinical needs and transform the landscape of cancer treatment. The expanding number of ongoing pipelines in FDA highlights the promise of ADCs as the new modality and component of combination regimens with ICB.

As is mentioned, the combination treatment with ICB and other modalities, such as chemotherapy, radiotherapy, ADCs or another ICB with different target antigen, has shown potentials in overcoming resistance and improving overall survival in various cancers^5,23,53^. Chemotherapy can have immunomodulatory effects by inducing immunogenic cell death and eliciting release damage-associated molecular patterns. Previous study also shows elevated intragenic rearrangement burden of patient and double-strand break DNA score after receiving platinum-based chemotherapy^27^. These effects make chemotherapy an appropriate partner for ICB, as the increased antigen release from dying tumor cells can alter the tumor microenvironment and thereby synergize with the immune activation caused by checkpoint blockades^54^. Besides chemotherapy, radiotherapy is also investigated in combination with ICB either before or after^53^. Radiation has been shown to increase the tumor lymphocyte infiltration, making the tumor more susceptible to immune attack. When combined with ICBs, radiation may help overcome resistance by increasing tumor immunogenicity. Early results also suggest that this approach may improve response rates and clinical outcomes in patients with limited response to either modality alone^55^.

## Conclusion

Immunotherapy by ICB has completely changed the cancer treatment. Despite its improved clinical outcome, only a small fraction of patient can benefit from the ICB treatment. As the field of immuno-oncology continues to evolve, the development of more robust biomarker and sophisticated treatment regimen will address such limitations of current ICB therapies, broaden the population of beneficiary patients and extend the duration of response. In this review, we discuss the biomarker discovery from three aspects that influence ICB efficacy: tumor genome, microenvironment and host immune profile. In addition, we also review the current transcriptomic curated databases for ICB biomarker studies and provide IOhub, a web portal and database that incorporates the largest number of bulk and single-cell datasets for responsive tumor samples to the ICB. Finally, we shed light on the use of bispecific antibodies and combination treatment in immunotherapy. With ongoing clinical trials and preclinical studies, these innovations are expected to improve response rates to the ICB, reduce cancer resistance, and ultimately enable precision immuno-oncology.

## Acknowledgement

This research was supported in part by The National Library of Medicine; National Cancer Institute at the National Institutes of Health grants R00LM013089, R01LM012011 and R01CA254274. The Department of Biomedical Informatics at University of Pittsburgh specifically disclaims responsibility for any analysis, interpretations or conclusions.

## Data Availability

The IOhub web portal is supported by the Center for Research Computing at University of Pittsburgh. The R Shiny App is accessible via https://shiny.crc.pitt.edu/iohub/. The processed normalized gene expression data are available upon request.

